# Analysis of patient data reveals novel cancer-relevant functions for GCN2/eIF2αK4

**DOI:** 10.1101/2025.03.18.643965

**Authors:** Vilte Stonyte, Christina Sæten Fjeldbo, Priyanka Swaminathan, Prabin Sharma Humagain, Laura Marian Valencia-Pesqueira, Lilian Lindbergsengen, Heidi Lyng, Beata Grallert

## Abstract

Numerous studies have shown that high GCN2 levels correlate with poor survival in a number of cancers. GCN2 has been known for over thirty years as a stress-response kinase, which phosphorylates the translation-initiation factor eIF2α, and thereby contributes to reprogramming of translation. Here we performed correlation analyses of GCN2 expression and that of other genes in a patient-derived sample set of cervical cancer samples. We found correlations not only with genes involved in stress responses, but also with genes involved in mitosis and cell migration. Our functional analyses confirmed that these correlations indeed reveal novel functions. Furthermore, our analyses of growth benefits associated with elevated GCN2 levels suggest that the novel functions can contribute to aggressive disease in cancers with high GCN2 levels.

## Introduction

All cells must be able to adapt to changes in their environment to survive in a dynamic environment. Stresses may arise from intrinsic sources (such as mistakes in protein folding, reactive oxygen species) or extrinsic sources (such as shortage of amino acids, hypoxia, damaging agents). One of the most important signaling pathways activated by these stresses is the integrated stress response (ISR), which culminates in reprogramming of translation to restore homoeostasis [1–4]. The ISR consists of several kinases which are each activated by their specific stress types, then phosphorylate the translation-initiation factor eIF2α. This leads to a reprogramming of translation, involving stimulation of translation of the mRNA encoding the transcription factor 4 (ATF4). ATF4, in turn, activates expression of genes involved in cell survival and recovery [5]. Dephosphorylation of eIF2α is thought to signal termination of the ISR and return to normal protein synthesis [6, 7].

Cancer cells are often surrounded by an unfavourable microenvironment, experiencing several kinds of stress, which they have to adapt to in order to successfully proliferate and metastasize. These stresses include endogenous stresses due to oncogene activation and increased translation, as well as a exogenous stresses such as a shortage of amino acids and oxygen in a growing tumour, both challenging the ISR [8–11]. The chronic ISR activation not only enables cancer cells to thrive in a hostile environment, but it can also be responsible for treatment resistance [12–14]. Given the importance of the ISR for cancer cells, small-molecule inhibitors to interfere with the ISR are actively developed for clinical application [10, 15, 16].

GCN2 is one of the ISR kinases, which has been known for over 30 years for its importance in the response to amino-acid starvation and other stresses, such as UV irradiation, proteotoxic stress and hypoxia [17, 18]. These stresses are highly relevant in the context of cancer. A number of studies revealed the importance of GCN2 in supporting tumor growth [11, 19–22], which is attributed to its role in stress responses. Recent reports also showed the involvement of GCN2 in shaping the immune response in the tumor microenvironment [23–25], raising interest in GCN2 as a therapeutic target. Furthermore, several recent studies reported activation of GCN2 by an off-target mechanism by other kinases, including agents already in clinical use or trials [26–28], again suggesting that interfering with GCN2 activity could improve the efficacy of other treatments.

However, there are some clues in the literature hinting that the canonical function might not be the only relevant one for cancer. First, a large-scale CRISPR-based study has shown that close to 13% of cancer cell lines (of ca. 2000) are dependent on GCN2 for survival, in contrast to <1% for each of the other three eIF2-α kinases [29, 30], suggesting that this sensitivity is not associated with the eIF2α kinase activity. Second, the four eIF2α kinases can compensate for each other at least in some model systems [11, 31], which is difficult to reconcile with the concept that the eIF2α kinase activity is the only reason for the importance of GCN2 in cancer. Third, GCN2 has been implicated in controlling ribosome biogenesis independently of its role in the ISR [32]. Fourth, GCN2 has been identified as essential for a cell-cycle checkpoint both in fission yeast and in budding yeast [33–35], pointing to functions in cell-cycle regulation. Furthermore, we recently reported that GCN2 is required for progression through mitosis in human cancer-derived cell lines, through regulation of PP1 activity, and independently of its role in the ISR [36].

To gain a better understanding of the importance of GCN2 in cancer, we turned to a biobank of cervical-cancer patient samples and explored correlations of GCN2 mRNA levels with that of other genes. Here we show that GCN2 mRNA levels correlate with expression levels of a number of genes, including, but not limited to, genes associated with stress responses. We also identified correlations with genes involved in mitosis, migration and cell motility, and immune responses. Deregulation of selected correlating genes involved in these pathways was also found in cell lines engineered to overexpress GCN2 or depleted for GCN2. Furthermore, we have performed functional studies to investigate the importance of GCN2 overexpression for mitosis and cell migration, functions predicted based on the analysis of patient data. We have shown that not only is GCN2 involved in both of these processes, but elevated levels of GCN2 confer advantages to cancer cells in the context of these functions.

## Results

### GCN2 expression levels correlate with that of genes involved in cancer-relevant processes

Consistent with the idea that GCN2 is important in cancer development, elevated GCN2 mRNA levels correlate with poor prognosis in several cancers [37, 38], including cervical cancer (Fig 1). To explore biological processes associated with GCN2, whole-genome gene-expression data of two cohorts of patients with cervical cancer were used (cohort 1, n = 156; cohort 2, n = 135). Spearman’s rank correlation analyses between the mRNA level of GCN2 and all other genes were performed, identifying 2629 and 1179 genes showing a significant (FDR < 0.05) negative and positive correlation with GCN2 in both cohorts, respectively (Table S1).

**Figure 1.**
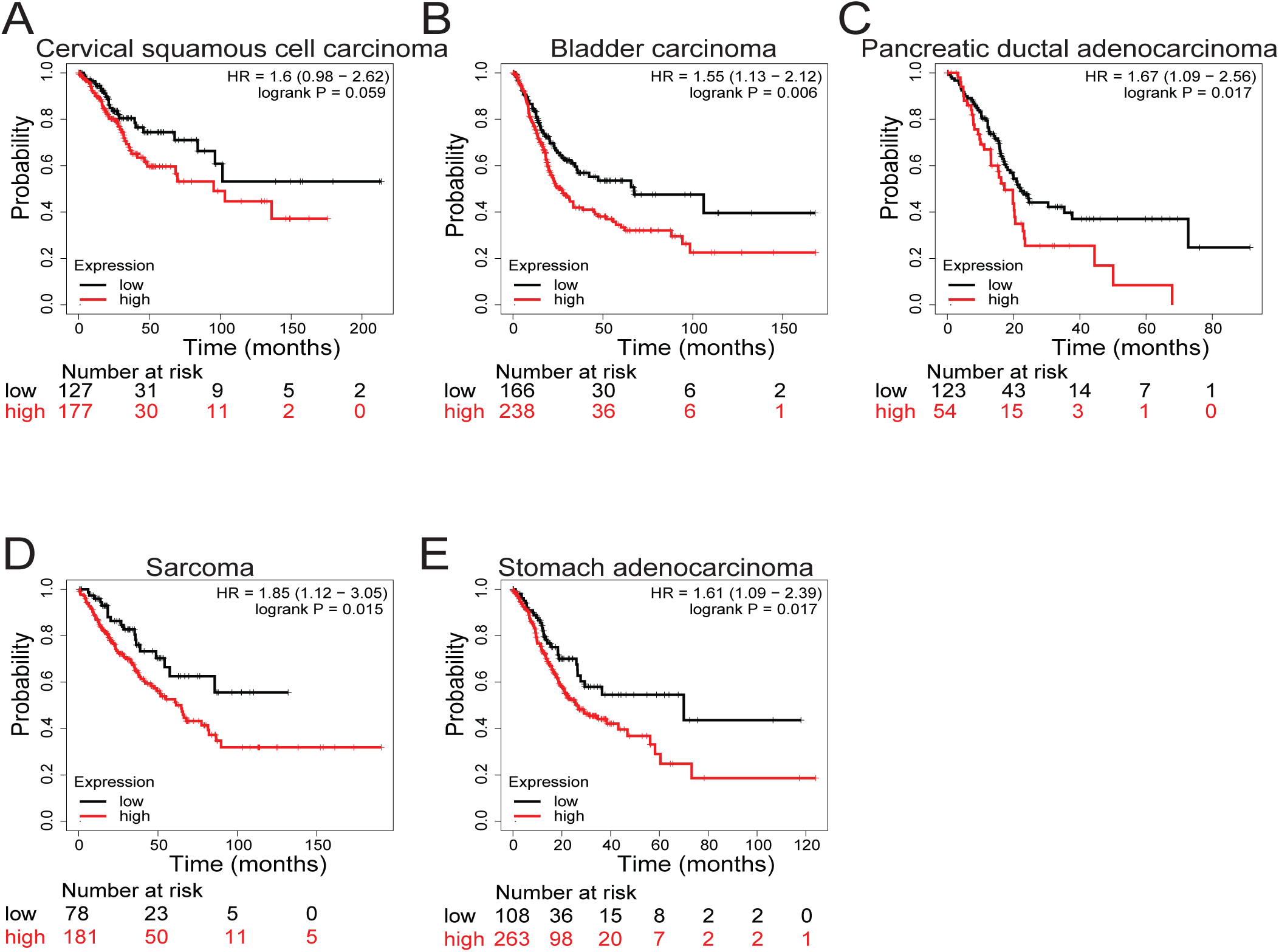
Elevated GCN2 expression levels correlate with poor prognosis in various cancers. Kaplan-Meier plots were generated in https://www.kmplot.com/analysis/ [37]

In order to validate our approach, we queried correlations we expected to find based on the canonical function of GCN2. GCN1 is thought to be required for GCN2 function in the ISR, associates with the ribosome and is involved in transmitting the signal upon starvation or other forms of stress leading to ribosome collisions to activate GCN2 [17, 39–43]. Given the close functional relationship, an overlap between genes correlating with GCN2 and GCN1 mRNA levels can be expected. Indeed, GCN2 levels significantly correlate with GCN1 levels in both cohorts (Fig S1A), consistent with their functional correlation. Furthermore, of 3798 genes correlating with GCN2 2231 (59%) also correlated with GCN1 (Fig S1B). As further validation, known ATF4 targets were searched for in the list of genes whose mRNA level correlates with that of GCN2 and/or GCN1. Of 109 previously identified target genes [11, 44] 40 genes were also found to correlate with GCN2 and/or GCN1 in the patient data set (Table S2). The identification of these expected correlations validates our approach.

Next, biological processes associated with the GCN2-correlating genes were predicted by subjecting the 1000 genes with the highest positive correlation and the 1000 genes with the lowest negative correlation to GSEA (Fig. 2A, TableS3). This analysis revealed biological processes previously associated with GCN2 and the integrated stress response; for example the Myc pathway [11, 45], TGF-β signaling [30], oxidative phosphorylation [46, 47], inflammation [23, 48–50], or DNA repair [51]. Intriguingly, the analysis also highlighted hallmarks not generally associated with GCN2, including “mitotic spindle”, “G2/M checkpoint”, and “apical junctions”, pointing to possible novel functions for GCN2.

**Figure 2.**
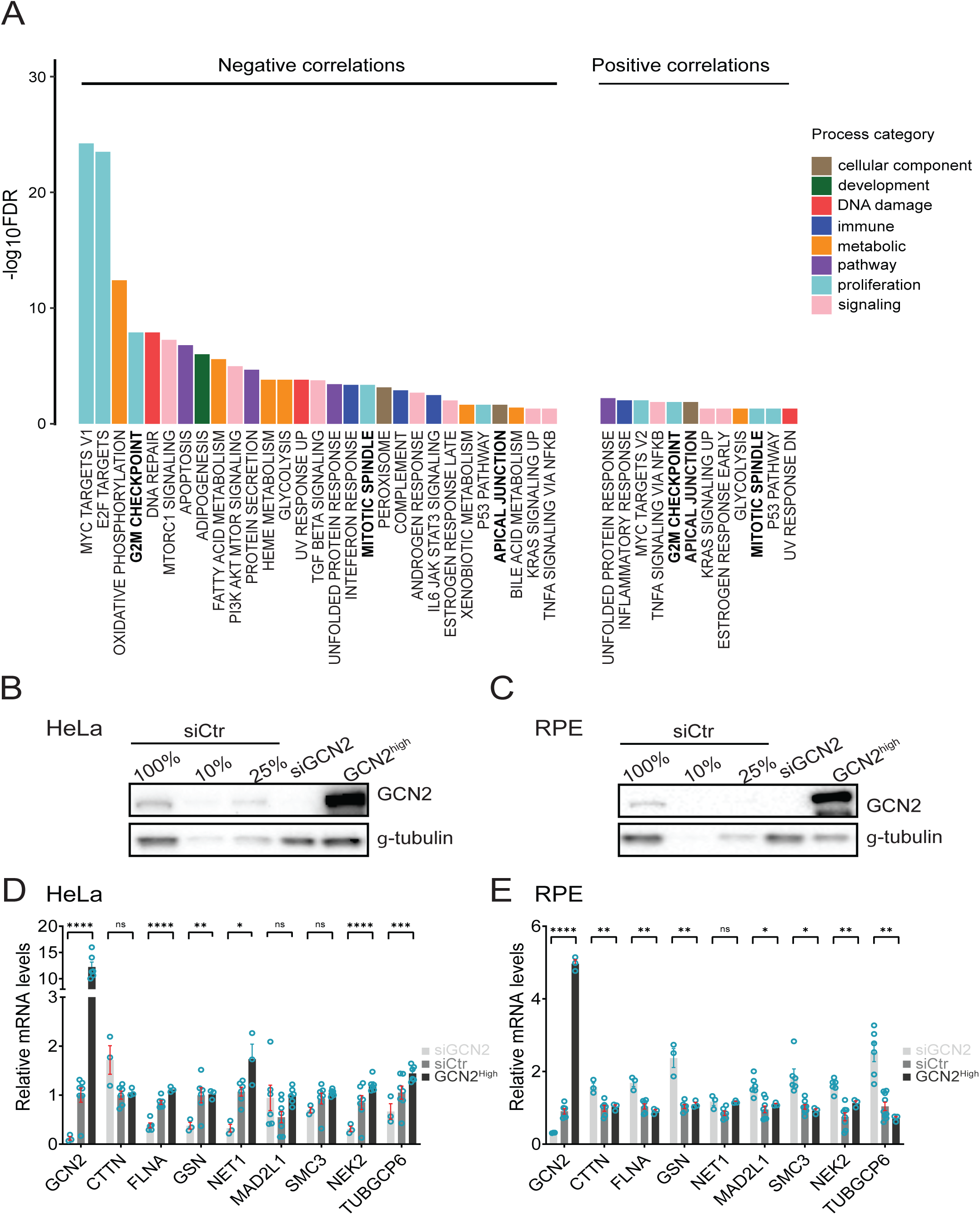
**A**: GSEA of GCN2-correlating genes. Enriched gene sets (FDR < 0.05) from the MSigDB Hallmark gene set for negatively (left) and positively (right) correlating genes. Selected gene sets indicating possible new functions of GCN2 pursued in this study are shown in bold. The results of the GSEA of the top-ranked 1000 correlating genes in each direction are shown in Table S3. **B, C** Representative immunoblots to show GCN2 levels in (**B**) HeLa and (**C**) hTert-RPE1 cell lines. Cells were transfected with GCN2-targeting siRNA-s to deplete (siGCN2) or were engineered to stably overexpress GCN2 by lentiviral transduction (GCN2^high^). To estimate the efficiency of depletion, different amounts (100%, 10% and 25%; corresponds to 30, 3 and 7.5 µg protein, respectively) of the lysate from cells transfected with control siRNA was loaded, along with 30 µg each of the transfected and the overexpressing samples. γ-tubulin is shown as a loading control. **D, E** mRNA levels of selected transcripts in (**D**) HeLa and (**E**) hTert-RPE1 cells. mRNA levels were normalized to TBP and to GCN2 levels in the parental cell line. Each gene was analyzed in at least three independent experiments. The statistical analyses (unpaired t-test, Benjamini, Krieger and Yekutieli method) compared values between the depleted and overexpressing samples, ****p<0.0001, (D) p=0.0879 for CTTN, **p=0.0019 for GSN, *p=0.0111 for NET1, p=0.8585 for MAD2L1, ***p=0.0002 for SMC3, ***p=0.0009 for TUBGCP6, (E) **p=0.0096 for CTTN, **p=0.0025 for FLNA, **p=0.0088 for GSN, p=0.9396 for NET1, *p=0.0128 for MAD2L1, *p=0.0149 for SMC3, **p=0.0037 for NEK2, **p=0.0020 for TUBGCP6.

To find more specific functions and identify key genes within the selected hallmarks, we analyzed GO-terms associated with the correlating genes overlapping with these hallmarks. Interestingly, this analysis revealed functions such as positive regulation of cell migration and cell motility, regulation of actin filament polymerization, as well as mitotic spindle assembly (Fig S2, Table S4). Furthermore, a comparison of the gene set correlating with GCN2 in the biobank against the MSigDB C5 GO BP database revealed 94 positively correlating genes with the GO term Locomotion and 87 of these are also associated with the GO term cell migration (data not shown).

To further probe the functional significance of the above findings, we selected representative genes correlating in the patient data in each of the pathways of interest, and investigated whether the mRNA levels of these genes also correlate in isogenic cell lines, where we either depleted GCN2 by siRNA (siGNC2) or overexpressed it (GCN2^high^). We used HeLa cells (of cervical-cancer origin) and a non-transformed epithelial cell line, hTert-RPE1 for these studies. GCN2 levels after depletion or overexpression were shown by immunoblotting (Fig 2 B, C). Intriguingly, the expression of several of the tested genes was altered when GCN2 was overexpressed or depleted (Fig 2D, E). However, the extent of these changes was very modest. Furthermore, neither the direction (up- or downregulation) nor the extent of the changes was consistent between the transformed and non-transformed cell lines.

To further explore the significance of the correlations found in the patient dataset, we turned to a large-scale database [29] to query correlations between GCN2 expression and expression of the selected genes identified as correlating in the patient data set. Remarkably, most of the correlations could be observed also in the large-scale dataset across many different cancers (Table 1). Furthermore, even when a correlation was not significant, a close functional relative, such as another component of the same complex or pathway, showed significant correlation (Table 1). These trends and correlations cannot be interpreted as direct evidence for GCN2 regulating the expression levels of the tested genes. However, the altered expression levels of several genes involved in the indicated pathways is consistent with a model where GCN2 is involved in the relevant processes. In this scenario, altered GCN2 levels have an impact on the given process, and thereby on the expression levels of genes involved in the pathway. Altogether, the observed correlations and expression patterns indicate novel functions of GCN2.

**Table 1.** Correlations of selected genes with GCN2 in DepMap.

| Significant correlations of selected genes in DepMap |  |  | Genes that correlate with GCN2 in patient data but not in DepMap, listed in line with functional relatives significantly correlating in DepMap. |  |  |
| --- | --- | --- | --- | --- | --- |
| Gene | Pearson | p-value | Gene | Pearson | p-value |
| **FLNA** | 0.447 | 7.07E-70 |  |  |  |
| **GSN** | 0.296 | 6.58E-30 |  |  |  |
| **NEK7** | 0.281 | 7.80E-27 | **NEK2** | 0.042 | 1.16E-01 |
| **CTTN** | 0.275 | 9.38E-26 |  |  |  |
| **MAD3L** | 0.154 | 5.97E-09 | **MAD2L1** | 0.032 | 2.26E-01 |
| **SMC2** | 0.131 | 3.99E-07 | **SMC3** | 0.061 | 1.81E-02 |
| **SMC5** | 0.125 | 1.36E-06 |  |  |  |
| **NET1** | -0.106 | 6.85E-05 |  |  |  |
| **TUBGCP6** | -0.091 | 6.40E-04 |  |  |  |
| n=1406 |  |  |  |  |  |

Genes identified as correlating in the biobank are shown in bold.

#### GCN2 in mitosis

G2/M checkpoint and mitotic spindle were hallmarks associated with several genes correlating with GCN2 mRNA levels, pointing to a function in mitosis. This finding further confirms and supports our recent report, where we identified PP1α and PP1γ as novel GCN2 substrates, and suggested that GCN2 affects the delicate balance of phosphorylation - dephosphorylation of numerous proteins controlling chromosome alignment and spindle function by regulating PP1α and PP1γ [36]. Remarkably, this function was not essential for normal mitotic progression in the non-transformed hTert-RPE1 cell line, but was essential in several cancer cell lines, including HeLa of cervical cancer origin [36]. We further investigated the impact of GCN2 inhibition on mitosis in other cervical-cancer-derived cell lines. SiHa and Caski cells also had a severe mitotic phenotype upon inhibition of GCN2 and could not align their chromosomes in a metaphase plate. Most HeLa cells eventually performed anaphase in spite of chromosome alignment defects and mitotic slippage was less penetrant [36].

Here we find that in SiHa and Caski cells fewer cells were able to perform anaphase and instead slipped mitosis (as judged from condensing and then decondensing their chromosomes) (Fig 3 A, B). Closer inspection revealed that the vast majority of cells undergoing “slippage” displayed spindle collapse, including multipolar spindles (Fig 3C, Movie 1-4), which was likely the reason for their failure to separate their chromosomes into two daughter nuclei. Notably, multipolar spindles were also observed in HeLa cells [36], even if this particular mitotic phenotype was less penetrant. While the exact outcome depends on the interplay of several regulatory mechanisms, which are deregulated in different ways and extents in different cancer cells, a common theme is that inhibiting GCN2 activity in cervical-cancer cells interferes with mitotic progression (Fig 3A, B and [36]) and leads to reduced cell viability (Fig S3A and [36]).

**Figure 3.**
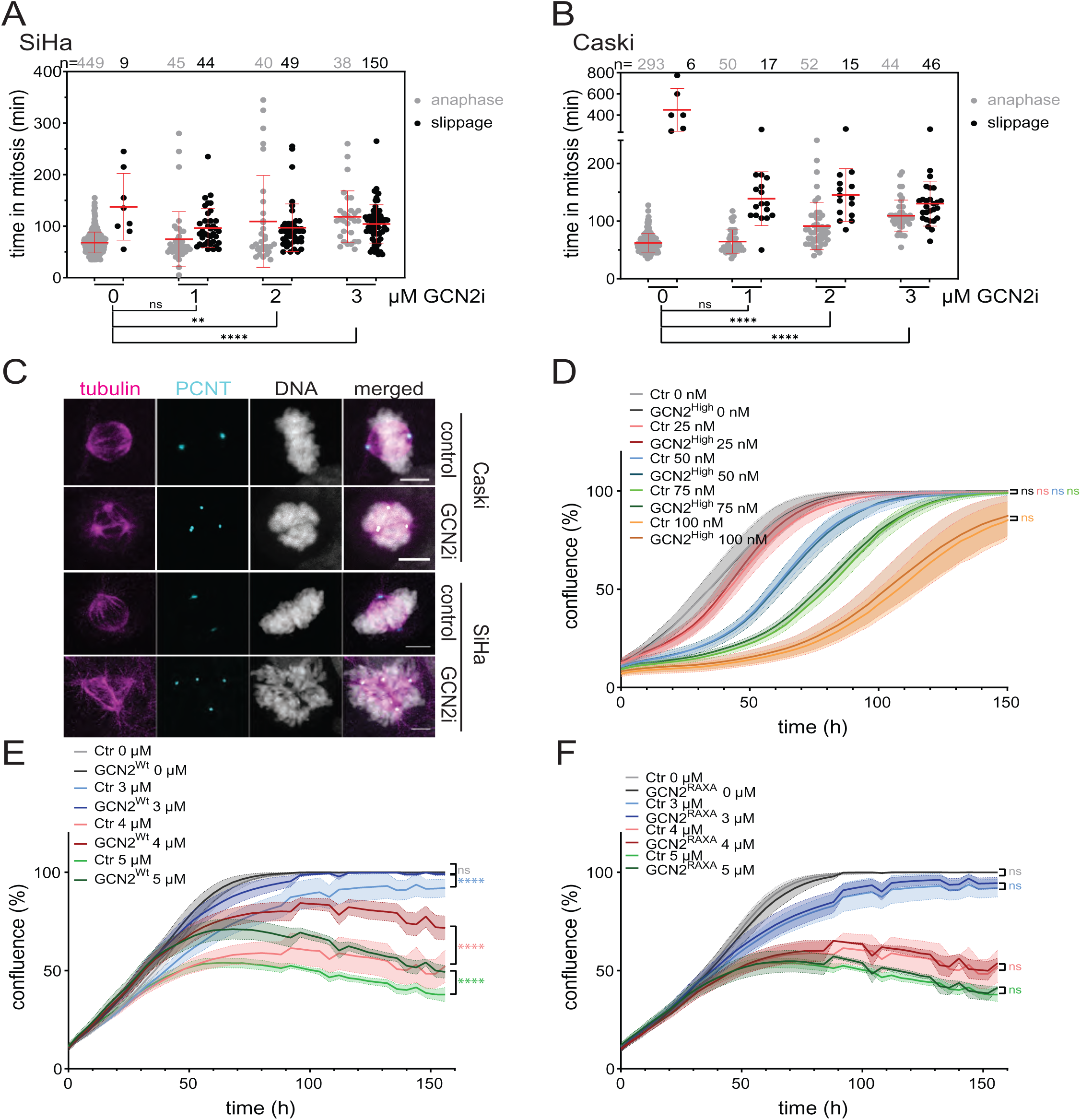
**A, B** Progression through mitosis in the presence of GCN2i at the indicated concentrations was observed by live-cell imaging in **A** SiHa and **B** Caski cells. The time in mitosis is shown for each cell. Prophase was judged by the first frame showing chromatin condensation), anaphase was scored based on chromosome separation and mitotic slippage was scored based on decondensation of the DNA in the absence of chromosome separation. Brown Forsythe and Welch Anova test was performed on the total time in mitosis. SiHa 0 µM compared to 1 µM p=0.1236; 2 µM *p=0.0014; 3 µM ****p<0.0001. Caski 0 µM compared to 1 µM p=0.0551; 2µM and 3 µM **** p<0.0001. **C** Caski and SiHa cells were incubated in the absence (control) or presence (GCN2i) of GCN2i for 8 hours and fixed for immunofluorescence. Representative images of mitotic cells stained for tubulin (magenta), pericentrin (PCNT, cyan) and DNA (grey) are shown. Scale bars represent 5 µm. **D** High levels of GCN2 give no protection in starved cells. Hela cells transduced to overexpress GCN2 (GCN2^high^) and the parental cell line (Ctr) were grown in the presence of HFG at the indicated concentrations and observed in Incucyte. Data shown are from three independent experiments. Mean ± SEM are shown, non-linear regression, p=0.9867 for 0 nM, p=0.1607 for 25 nM, p=0.9012 for 50 nM, p=0.1994 for 75 nM, p=0.3283 for 100 nM. **E** High levels of GCN2 gives protection in cells exposed to MPS1i. Hela cells transduced to overexpress GCN2 (GCN2^WT^) and the parental cell line (Ctr) were grown in the presence of MPS1i at the indicated concentrations and observed in Incucyte. Data shown are from four independent experiments. Mean ± SEM are shown, non-linear regression, p=0.3393 for 0 µM, ****p<0.0001 for 3 µM, 4 µM, and 5 µM. **F** Protection in cells exposed to MPS1i is lost in cells expressing GCN2^RAXA^. HeLa cells transduced to express siRNA-resistant GCN2^RAXA^ were transfected with GCN2-targeting siRNA (GCN2^RAXA^), grown in the presence of MPS1i at the indicated concentrations, and observed in Incucyte. Data shown are from four independent experiments. Mean and SEM are shown, non-linear regression, p= 0.5069 for 0 µM, p= 0.1346 for 3 µM, p=0.4104 for 4 µM, p=0.4254 for 5 µM. Note that the data shown for control cells are the same as those shown in Figure 3E.

The involvement of GCN2 in mitosis raises the question whether it is important in the context of the correlation of high GCN2 levels with poor survival. To test the impact of high GCN2 levels on cell growth under stress, we considered both the canonical role under amino-acid starvation and the mitotic role. To test whether high levels of GCN2 confer growth advantages under starvation conditions, we grew HeLa cells and GCN2^high^ HeLa cells in the presence of halofuginone (HFG), an inhibitor of glutamyl-prolyl tRNA synthetase, and monitored cell growth. Surprisingly, high levels of GCN2 did not confer a growth advantage in face of starvation (Fig 3D). In light of the novel function in mitosis, a plausible hypothesis is that high levels of GCN2 can protect the cells from some forms of mitotic stress. To test this hypothesis, we exposed HeLa and GCN2^high^ HeLa cells to inhibitors affecting mitotic processes. When cells were exposed to an inhibitor of the SAC kinase MPS1, high GCN2 levels led to a significantly improved survival (Fig 3E, GCN2^WT^). Notably, overexpression of a GCN2^RAXA^ mutant that cannot bind PP1 [36] did not confer this growth advantage in the presence of the MPS1 inhibitor (Fig 3F), confirming that this effect is due to GCN2’s mitotic function though PP1 regulation. A similar effect was observed using STLC, an inhibitor of the mitotic kinesin Eg5 / KIF11 (Fig S3B, C), further supporting the notion that high levels of GCN2 confer growth advantages in the face of some kinds of mitotic stress in cancer cells.

#### GCN2 affects cell migration

Another intriguing pathway associated with GCN2 in our analyses was cell migration and motility, which prompted us to investigate a role for GCN2 in this process. mRNA levels encoding several actin remodelers (CTTN, FLNA, GSN) showed significant correlationsin the cervical cancer biobank (Table S1), in the DepMap database (Table 1) and in qPCR analyses in the selected cell lines (Fig 2D, E).

To address a role for GCN2 in cell migration, we assessed the cells’ ability to move towards a chemoattractant in a transwell migration assay, using FBS as a chemoattractant, employing isogenic cell lines overexpressing (GCN2^high^) or depleted for GCN2 (siGNC2). The capacity for directional migration correlated with GCN2 levels: not only did cells have a reduced capacity when depleted for GCN2, but, remarkably, cells overexpressing GCN2 had an increased capacity for directional migration (Fig 4A,B).

**Figure 4.**
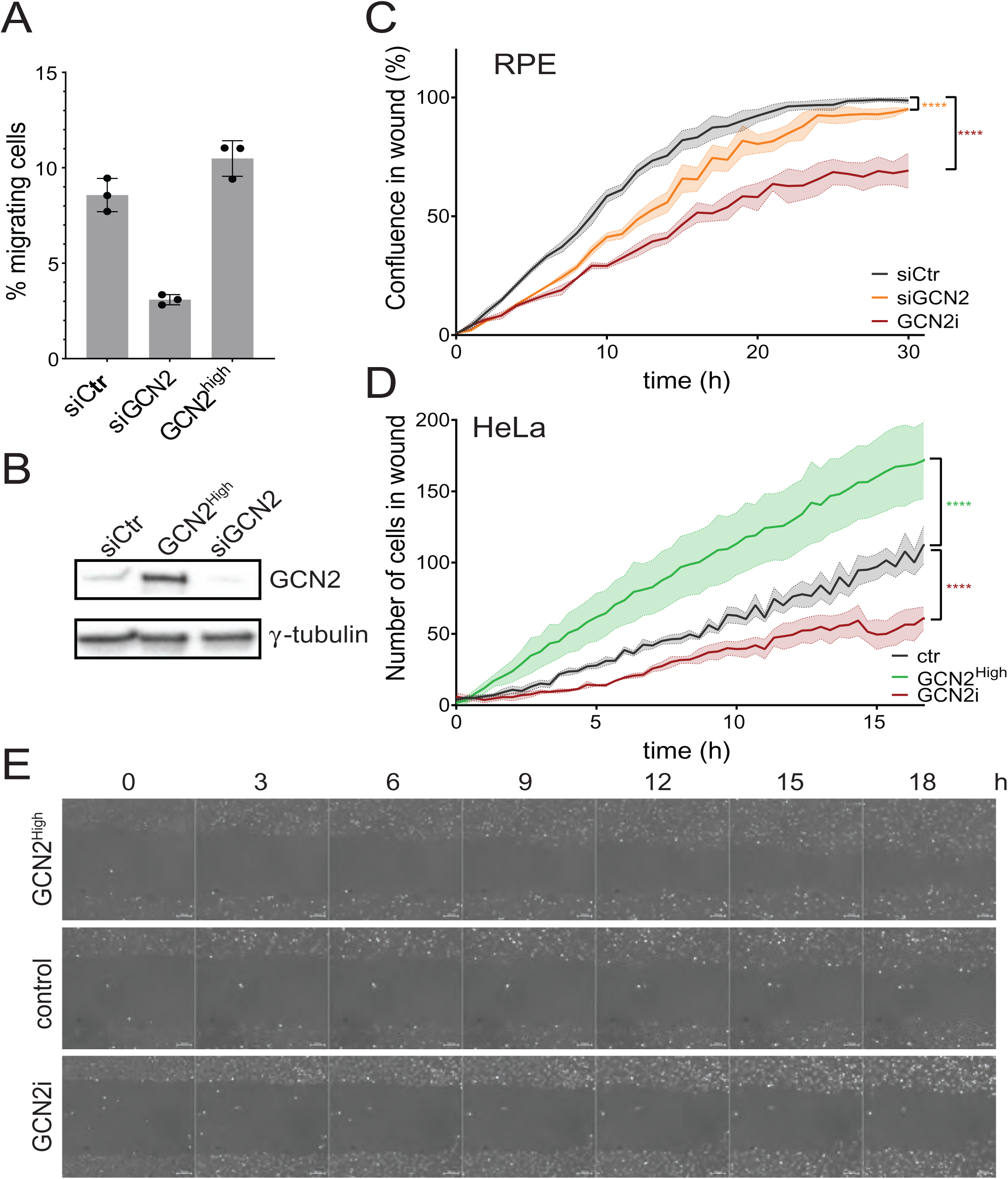
**A** Migration capacity correlates with GCN2 levels in hTert-RPE1 cells. hTert-RPE1 cells were transfected with GCN2-targeting siRNA for 48 h (siGCN2) or transduced to stably overexpress GCN2 (GCN2^high^). Cells were seeded without FBS into transwell chambers with 8 µM pore size and exposed to an FBS gradient for 16 hours. Migrated cells were counted after DAPI staining and normalized to seeding controls. Three independent experiments, mean and STDEV are shown. **B** Immunoblots of lysates of cells from the experiment shown in A. γ-tubulin is used as a loading control. **C** hTert-RPE1 cells were transfected with GCN2-targeting siRNA (siGCN2) or treated with 2 µM GCN2i (GCN2i). Cells were grown to confluence and wounds were created using a Wound Maker tool (Sartorius) and healing was monitored in Incucyte. Three (siGCN2) or five (GCN2i) independent experiments, mean ± SEM are shown, non-linear regression, ****p<0.0001 **D** HeLa cells treated with 2 µM GCN2i (GCN2i) or transduced to stably overexpress GCN2 (GCN2^high^) were seeded into Ibidi culture inserts and grown to confluence. After removal of the inserts the cells were observed by live-cell imaging. The number of cells migrating into the initial wound area is shown. Three (GCN2i) or four (GCN2^High^) independent experiments, mean and SEM are shown, non-linear regression, ****p<0.0001 **E** Representative images from an experiment shown in (D).

To further explore the relevance of GCN2 function for cell migration, we performed wound-healing assays using hTert-RPE1, HeLa, and Caski cells. In the absence of GCN2 function migration into the gaps was severely impaired in all three cell lines (Fig 4C, D, Fig S3D), consistent with a previous observation using keratinocytes [52]. Notably, GCN2 is not essential for mitosis in RPE cells [36], further supporting the notion that the observed impaired wound healing is not a result of impaired cell division but of impaired cell migration. Furthermore, HeLa cells engineered to overexpress GCN2 (GCN2^high^) were able to close the gap faster than control cells (Fig 4D, E, Movie s 5-7), suggesting that high levels of GCN2 in cancer cells lead to higher capacity to migrate. This feature might promote the ability to metastasize, consistent with the higher mortality rate correlating with high GCN2 levels in patients.

Directional cell movement requires perceiving and interpreting spatial signals, as well as rearranging the cytoskeleton to achieve movement. In order to address whether the loss of ability to respond to spatial cues is the reason for the impaired directional movement, we used a chemotaxis assay where cells were seeded in medium without FBS, challenged with an FBS gradient, and observed by live-cell imaging. Consistent with the results of the wound-healing and transwell assays, in the absence of GCN2 activity both HeLa and Caski cells had reduced ability for chemotactic movement (Fig 5A, B). Remarkably, the speed of movement was considerably impaired in the absence of GCN2 function in both cell lines, (Fig 5C, E), while the directness of movement was not much impaired (Fig 5D, F). Furthermore, overexpression of GCN2 in HeLa cells, which led to faster wound healing (Fig 4D, E), correlated with increased cell velocity (Fig 5G) rather than with increased directness of movement (Fig 5H). These results suggest that the primary reason for poor wound healing or chemotactic movement in the absence of GCN2 is not an impaired ability to respond to spatial cues, but rather impaired movement, and faster wound healing in cells with elevated GCN2 levels correlates with faster movement. To further explore this conclusion, we also assessed cell velocity under normal growth conditions where the cells were not exposed to any gradient or wound. Consistently with the findings of the chemotaxis assays, cell movement was affected by GCN2 inhibition in both HeLa and RPE cells and cells overexpressing GCN2 had a higher velocity (Fig S3D). These data strongly suggest that GCN2 affects directional cell migration through affecting cell movement, although a smaller impact on directness of movement cannot be excluded.

**Figure 5.**
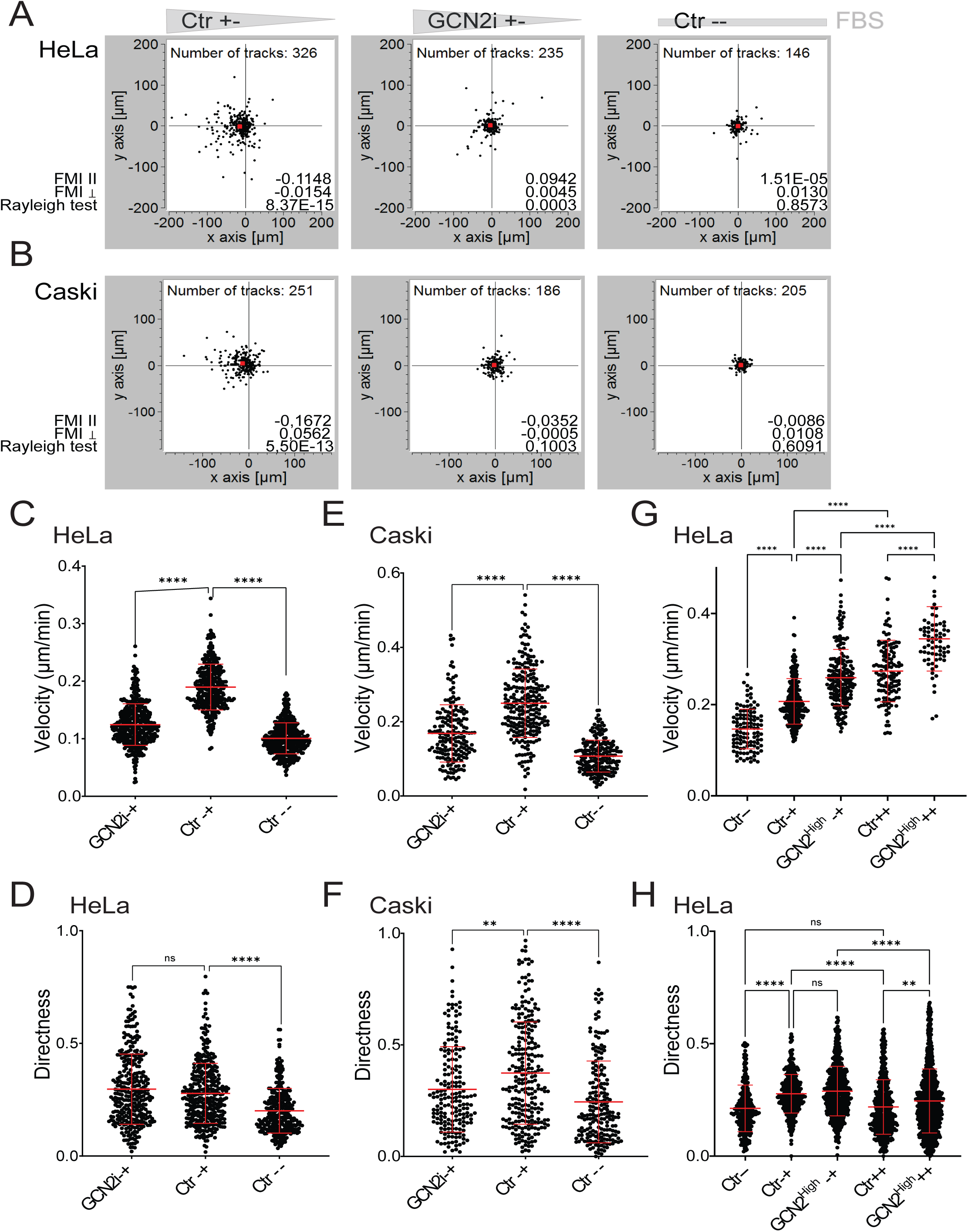
**A, B** HeLa (**A**) or Caski (**B**) cells were seeded into Ibidi µ-Slide Chemotaxis with (+-) or without (--) an FBS gradient and observed by live-cell imaging. Shown are endpoints of tracks from representative experiments. Red squares indicate the center of mass displacement. +- denotes the presence of FBS gradient, -- denotes no FBS on either side; the concentration of FBS is also indicated by the grey shaded triangles. **C, E, G** Velocity of the indicated cell lines and treatments was determined in chemotaxis experiments in three (**C,G**) or two (**E**) independent experiments. -+ denotes the presence of FBS gradient, -- denotes no FBS on either side. Cumulative results are shown. Welch’s two-tailed t test, ****p<0.0001 **D, F, H** Directness of movement of the indicated cell lines and treatments was determined in chemotaxis experiments in three (**D, H**) or two (**F**) independent experiments. -+ denotes the presence of FBS gradient, -- denotes no FBS on either side. Cumulative results are shown. Welch’s two-tailed t test, p=0.0963 for GCN2i versus Ctr -+; ****p<0.0001 for Ctr -+ versus Ctr --.

Previous studies implicated the canonical function of GCN2 in wound healing [47, 52]. In order to explore whether eIF2α is the only relevant substrate in this function, we made use of ISRIB, a drug that binds elF2B and thereby antagonizes the inhibitory effect of phosphorylated eIF2α [53] and compared its effects on wound healing to that of GCN2 inhibition. ISRIB impaired the wound-healing capacity of the cells, consistent with an involvement of eIF2α phosphorylation and the results of the previous reports. Interestingly, GCN2 inhibition had a more severe effect in RPE, HeLa (Fig 6A, B, Fig S4D) and Caski (Fig S3E) cells than treating the cells with ISRIB, using drug concentrations that abolish the starvation-induced induction of GADD34 (Fig S4A, B). These results suggest that additional substrate(s) are important for this function. We recently identified PP1α and γ as novel GCN2 substrates and showed that a GCN2^RAXA^ mutant, unable to bind PP1 PP1α and γ, is still able to phosphorylate eIF2α [36]. To address the importance of PP1 binding we employed stable cell lines depleted for the endogenous GCN2 by siRNA, and expressing siRNA-resistant wild-type (GCN2^WT^), kinase-dead (GCN2^K619R^) or RAXA-mutant GCN2 (GCN2^RAXA^). Neither mutant could rescue the migration defect seen in the absence of GCN2 in (Fig 6C, D, E, F), and the effect of ISRIB was additive with the effect of the RAXA mutation (Fig 6E, F). These data suggest that eIF2α, and PP1α and/or PP1γ are both relevant for the function of GCN2 in migration.

**Figure 6.**
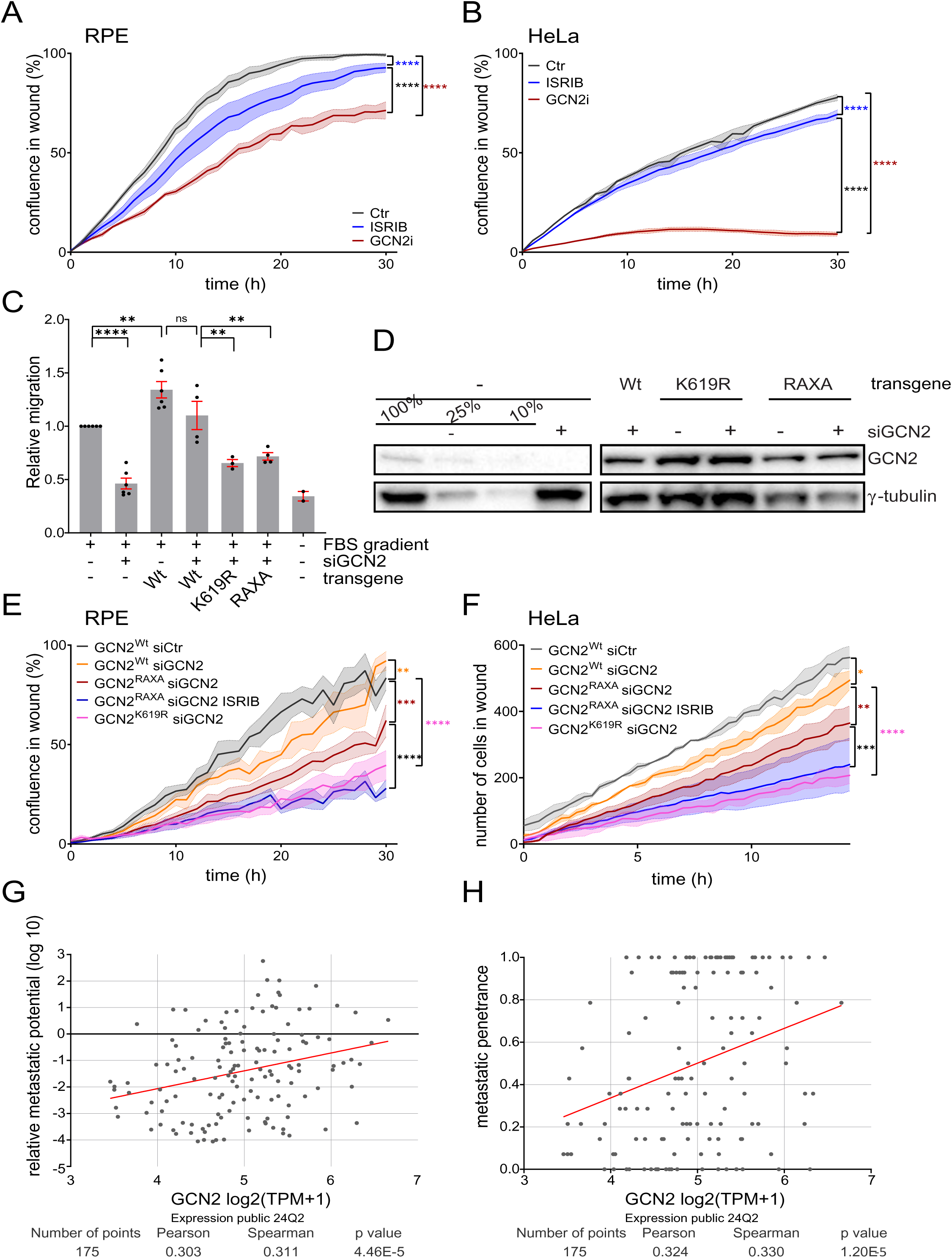
**A**, hTert-RPE1 and **B,** HeLa cells were grown to confluence and wounds were created using a Wound Maker tool (Sartorius) and healing was monitored in Incucyte. ISRIB or GCN2i was added to 200 nM or 2 µM, respectively. Four (hTert-RPE) or three (HeLa) independent experiments, mean and SEM are shown, non-linear regression, ****p<0.0001 **C** hTert-RPE1 cells transduced with GCN2-siR carrying the indicated mutations were transfected with control or GCN2-targeting siRNA for 48 h. Cells were seeded without FBS into transwell chambers with 8 µM pore size and exposed to an FBS gradient for 16 hours. Migrated cells were counted after DAPI staining and normalized to seeding controls, then to migration in cells transfected with control siRNA. Mean ± SEM are shown, results from at least three independent experiments. One-way Anova,****p<0.0001 siCtr versus siGCN2; **p=0.0022 siCtr versus GCN2^Wt^ siCtr; **p=0.0051 GCN2^Wt^ siGCN2 versus GCN2^RAXA^ siGCN2; **p=0.0025 GCN2^Wt^ siGCN2 versus GCN2^K619R^ siGCN2 **D** Knock-down efficiency and expression of the mutant transgenes in the experiments shown in C was tested by immunoblotting. γ-tubulin is shown as loading control. **E** hTert-RPE1 cells transduced with doxycycline-inducible siRNA-resistant GCN2 carrying the indicated mutations were transfected with control (siCtr) or GCN2-targeting siRNA (siGCN2) and grown to confluence in the presence of doxycycline. Wounds were created using a Wound Maker tool (Sartorius) and healing was monitored in Incucyte. ISRIB was added to 200 nM after wounding. Averages (lines) and SEM (bands) from six independent experiments are shown. Non-linear regression, **p=0.0028, ***p=0.0002, **** p<0.0001. **F** HeLa cells transduced with doxycycline-inducible siRNA-resistant GCN2 carrying the indicated mutations were transfected with control (siCtr) or GCN2-targeting siRNA (siGCN2), seeded into Ibidi culture inserts, and grown to confluence in the presence of doxycycline. After removal of the inserts the cells were observed by live-cell imaging. ISRIB was added to 200 nM after removal of the inserts. The number of cells migrating into the initial wound area is shown. Averages (lines) and SEM (bands) from three independent experiments are shown. Non-linear regression, ****p<0.0001, *p=0.0113, **p=0.0072, ***p=0.0006 **G, H** Metastatic potential (G) and penetrance (H) in metastatic cancers as a function of GCN2 mRNA levels. Data were downloaded from depmap.org and are based on a study by Jin et al (2020), which reported metastatic-potential profiling of ca. 500 human cancer cell lines derived from 21 types of solid tumour in immunodeficient murine models [54]. Metastatic potential is calculated based on the mean cancer-cell numbers detected in the target organs. Metastatic penetrance refers to the proportion of mice displaying metastases with the given cell line [54].

Regardless of the exact mechanism, the impact of high GCN2 levels on wound healing and migration capacity of the cells (Fig 4-6) suggests that elevated GCN2 levels might be beneficial for cancer cells because of their increased capacity to move to other sites and metastasize. To further explore the idea that metastatic potential correlates with elevated GCN2 level, we queried the Metastasis Map (MetMap) database, which describes organ-specific metastasis patterns of ca. 500 human cancer cell lines in immunodeficient murine models [54]. In this sample set GCN2 expression levels clearly correlate with increased metastatic potential (Fig 6G) and penetrance (Fig 6H).

## Discussion

GCN2 has long been seen as an attractive target in the context of cancer. High levels of GCN2 in several cancers correlate with poor prognosis, as apparent in the depmap pan-cancer database [37, 38], as well as a number of previous studies. For example, Ye et al. [21] observed increases in both the total amount of GCN2 and phosphorylated GCN2 in colon, lung, breast and liver cancer tissue samples when compared with healthy tissue. Wang et al. [20] made similar observations in human oral squamous cell carcinoma, and, most recently, Furnish et al. [55] reported similar results in a prostate-cancer study. In addition, increased GCN2 expression levels in human papillary renal cell carcinoma were a powerful indicator of poor patient outcomes [56]. Furthermore, a recent study using the TCGA data revealed that GCN2 expression is elevated in 26 out of 29 cancer types [57].

However, elevated GCN2 expression levels do not always correlate with poor survival [37, 57]. Since exposure to stress is a general feature of cancers, this observation is difficult to interpret in the context of the canonical role in stress responses alone, indicating that additional functions also have to be considered.

In the current study, we performed correlation analyses of mRNA levels of GCN2 and that of other genes in patient samples from cervical cancer, and found correlations not only with stress-response genes, but also with genes involved in mitosis and cell migration. Our functional analyses confirmed that these correlations indeed reveal novel functions. Furthermore, our analyses of growth benefits associated with elevated GCN2 levels suggest that the novel functions can contribute to aggressive disease in cancers with high GCN2 levels.

The mitotic role suggested from the analysis of patient data was confirmed in several cervical cancer cell lines. While the phenotype was not exactly the same in different cell lines, a common theme was that GCN2 inhibition severely interferes with mitotic progression in each of the cervical cancer cell lines we tested, as previously reported for cell lines derived from other types of cancer [36]. These data further support our previously published findings [36] and highlight the relevance of GCN2’s mitotic role for cancer. Furthermore, the requirement for GCN2 for faithful mitosis in cancer cells is one plausible reason for why many cancer cell lines are dependent on GCN2 for growth in the absence of stress. The finding that mitosis in cancer cells is particularly vulnerable to GCN2 inhibition highlights the potential to use GCN2 inhibitors as a highly cancer-selective antimitotic strategy. Furthermore, we have shown that cells engineered to overexpress GCN2 were resistant to some forms of mitotic stress. This might be an important factor contributing to the poor prognosis of patients with cancers with high levels of GCN2. Cancer cells often undergo tetraploidization and up to 70 % of cancers are aneuploid, presumably providing growth advantages to the cancer cells [58–62]. However, in such cells aberrant and stressed mitoses are prevalent, leading to further genomic and choromosome abnormalities and further aberrant mitoses. In such a vicious circle even a small protection, as observed in cell lines with high GCN2 levels and exposed to drugs inducing mitotic stress in this study, might be beneficial for cancer cells. Furthermore, antimitotic agents are broadly used in chemotherapy and even if the protection by high levels of GCN2 in cell lines is modest, elevated GCN2 levels could contribute to resistance and should be considered when choosing therapeutic agents targeting mitosis.

The role in cell migration was confirmed in non-transformed epithelial cells, as well as two different cervical-cancer-derived cell lines. Previous studies implicated GCN2 in wound healing in keratinocytes [52] and poor wound healing in diabetes was associated with downregulation of GCN2 activity [47].

These studies attribute the involvement in wound healing to the canonical function through eIF2α phosphorylation. We previously identified PP1α and γ as novel GCN2 substrates in mitosis [36] and here investigated whether these substrates are relevant for cell migration. Several pieces of evidence suggest that eIF2α is not the only relevant substrate in the context of cell migration. First, in the present study GCN2 inhibition had a larger impact on cell migration than ISRIB, which counteracts eIF2α phosphorylation. Second, we previously showed that the two functions can be genetically separated by using a RAXA mutant, which is unable to bind PP1 but is able to phosphorylate eIF2α [36]. Here we found that the RAXA mutant was not able to rescue the migration defect after GCN2 knock-down. Third, the impact of ISRIB and the RAXA mutant on the wound-healing capacity was additive. Collectively, these data suggest that both substrates are important in the context of cell migration. The exact mechanism and affected PP1 substrates will be the subject of future studies.

Strikingly, cells engineered to overexpress GCN2 displayed an increased ability to migrate towards a chemoattractant compared to isogenic control cells with “normal” GCN2 levels, which is apparent in wound healing, transwell and chemotaxis assays. Directional movement requires that cells can rearrange their cytoskeleton to move, as well as to respond to directional cues. A previous study suggested that the main reason for the impaired wound healing when GCN2 is inhibited is a diminished localized production of ROS in the leading edge of the wound [52]. Consistent with this model, in our chemotaxis assays GCN2 inhibition did impair directional movement, even if it did not lead to a complete loss of directionality (Fig 5A). However, velocity was dramatically altered both in the presence of a GCN2 inhibitor and in cells overexpressing GCN2 (Fig 5), suggesting that loss of ability to move is the primary reason for the impaired wound healing. One plausible explanation for this effect is that GCN2 impinges on cellular metabolism, which in turn affects cell migration. A recent work reported that GCN2 deficiency in diabetic patients enhanced oxidative phosphorylation, thereby promoting macrophage polarization, which in turn play important regulatory roles in inflammation and tissue repair during wound healing [47]. An additional plausible model emerging from our results is that GCN2 can also affect cell movement through regulating the rearrangement of the cytoskeleton, possibly through modulating PP1 activity. Extensive further work will be required to explore this hypothesis.

Cancer cells expressing high levels of GCN2 display faster migration and chemotactic ability, which can contribute to a higher potential to metastasize. Consistently, higher GCN2 levels correlate with higher metastatic potential in the MetMap database. An increased capacity to metastasize could be an important factor determining poor prognosis for patients with cancers with elevated GCN2 levels.

The analysis of correlations in the patient data revealed important new functions in mitosis and cell migration, which are highly relevant for cancer progression and can at least in part explain the poor prognosis associated with high GCN2 levels in many cancers. These roles could make GCN2 a valuable target for cancer therapies, especially in cancers with high GCN2 expression and poor prognosis. The novel results presented here will aid in better understanding of the impact of interfering with GCN2 activity and in exploiting it in therapy.

## Acknowledgements

This work was supported by the Norwegian Research Council (#300288, BG), the Nansen Fond (University of Oslo), the Anders Jahres Fond (University of Oslo), and the Norwegian Cancer Society (#273730, BG).

We wish to thank the staff at the Light Microscopy Core Facility at the Institute for Cancer Research for advice and assistance with live-cell imaging and Mélanie Langiu for helpful comments on the manuscript.

## Methods

### Gene expression data from clinical samples of cervical cancer

Gene expression data of 291 locally advanced cervical cancer patients treated with curative chemoradiotherapy were used. Patient characteristics, treatment, tumor biopsies and data acquisition have been described previously [63, 64]. Whole genome gene expression data were generated by Illumina Bead Arrays, using total RNA extracted from pretreatment tumor biopsies. Log2-transformed data were used in the analyses. The patients were divided into two cohorts based on the Illumina Bead array version they were assayed by, i.e. WG-6 v3 (cohort 1, n = 156) or HT-12 v4 (cohort 2, n = 135). An estimate of the background level was calculated based on probes measuring Y chromosome genes [65], and probes where at least 95% of the 291 samples had a signal below this background estimate were excluded. Further, analyses were performed using probes present on both array versions, and with an Entrez gene id annotation, yielding 19811 probes measuring 14638 unique genes.

#### Identification of genes correlating with GCN2/GCN1

Spearman’s rank correlation analyses of GCN2 or GCN1 gene expression level versus the expression level of all other probes were performed for cohort 1 and 2 separately. The final list of correlating genes was generated as follows: Probes with significant (FDR <0.05) positive correlation in both cohorts, or negative correlation in both cohorts were included. Genes measured by more than one probe were kept only if all significantly correlating probes correlated with the same direction (i.e. positively or negatively correlating genes). The probe with largest average correlation coefficient (absolute value) for cohort 1 and 2 per gene was kept for the final list of correlating genes. This average correlation coefficient was also used for ranking the correlating genes for each direction.

#### Gene set enrichment analysis (GSEA)

Enrichment of the correlating genes in specific biological processes was searched for in the Molecular Signatures Database [66](MSigDB, v.7.4) Hallmark [67] and gene ontology gene set collectionshttps://tnmplot.com/analysis/. For the list of GCN2-correlating genes (i.e., the 1000 most correlating genes per direction based on the average correlation coefficient in cohort 1 and 2), the genes overlapping with the gene sets were identified, and statistical significance of the overlap was estimated using hypergeometric test.

All analyses of data from clinical samples were performed in R version 4.3.2 [68]. Correction for multiple testing in the spearman correlation analyses and GSEA, was performed according to the Benjamini and Hochberg procedure (FDR).

#### Cell culture

The human cervical-cancer cell lines HeLa, Caski, SiHa and LentiX cells were cultivated in DMEM (Dulbecco’s Modified Eagle’s Medium) (Invitrogen) supplemented with 10% fetal bovine serum (FBS) (Gibco) and 1% Penicillin/Streptomycin (P/S) (Gibco). The non-transformed epithelial cell line hTert- RPE1 was cultivated in DMEM/F12 (Invitrogen) supplemented with 10% FBS, 1% P/S and 0.01mg/ml Hygromycin B (Sigma). All cell lines were tested negative for mycoplasma contamination and grown at 37°C in a humidified environment with 20% O_2_ and 5% CO_2_.

Inhibitors were GCN2i: GCN2-IN-1 (MedChemExpress #HY-100877), Mps1i (MedChemExpress #HY- 14710), S-Trityl-L-cysteine (STLC) (Sigma #164739), ISRIB (MedChemExpress # HY-12495)

#### Immunoblots

Samples for immunoblots of mammalian proteins were prepared by one wash with cold 1× PBS and kept at -80°C until 2× Laemmli buffer was added to make whole cell lysates. Alternatively, cells were lysed with lysis buffer (100 mM NaCl, 50 mM Tris (pH 7.5), 2 mM MgCl_2_, 0.5% Triton X-100, 100 nM Calyculin A (LC laboratories), protease inhibitor cocktail (Roche) containing 100 U/ml benzonase (Merck), lysates were cleared by centrifugation at 13 000 rpm for 10 min and mixed with 4 x Laemmli buffer. ECL kits SuperSignal™ Western Blot Substrate Pico PLUS, Femto or Atto were used for detection. Images were captured with the Biorad Chemidoc system, and quantified using ImageLab software. Rabbit Gcn2T899-P 1:500, (R&D Systems, #AF7605); rabbit Gcn2, 1:1000 (Cell Signaling Technology, #3302); rabbit eIF2α-P, 1:1000 (Invitrogen, #44-728G); rabbit GADD34, 1:1000 (Proteintech, #10449-1-AP); rabbit ATF4, 1:500 (Cell Signaling Technology, #11815), rabbit eIF2α, 1:2000 (SCBT, #sc-11386); rabbit GAPDH 1:1000 (Cell Signaling Technology, #5174); mouse γ-tubulin 1:25000 (Sigma, #T6557).

#### Live-cell imaging

Cells were seeded into chambered coverslips (Ibidi, #80806) for analysis of mitotic progression, or into culture inserts (Ibidi, #80366) for wound healing assays. Cells were imaged in Fluorobrite DMEM medium supplemented with 10% fetal bovine serum, 1% penicillin-streptomycin, and Glutamax (ThermoFisher). SPY-DNA-650 (10[tnM; SpiroChrome) was used to visualize the nuclei, and SPY555- tubulin (1:5000, Spirochrome) was used to visualize tubulin.

Cells were imaged using a Nikon ECLIPSE Ti2-E inverted microscope (Nikon Corp, Tokyo, Japan) equipped with two Prime BSI sCMOS cameras (Teledyne Photometrics, Tucson, AZ, US), with a CFI Plan Apo l 40x (0.95 numerical aperture) (Fig3A, B) or a Plan Apo l 10x (0.45 numerical aperture) (Fig4D, 5, 6F, S3D, E, S4) objective. Alternatively, a DeltaVision Elite widefield microscope (Applied Precision), equipped with a live cell Elite TruLight Illumination System and a CoolSNAP HQ2-ICX285 camera, using an Olympus UPlan SApo 20× (0.75 numerical aperture) objective.

Time-lapse images (10 z-sections 0.8 µm apart) were acquired every 5 minutes (Fig3A, B), or every 20-30 minutes for wound-healing and chemotaxis assays. Anaphase was scored based on chromosome separation. The microscope stage was kept at 37°C by a temperature-controlled incubation chamber during live observation.

Images obtained with the Nikon ECLIPSE Ti2-E were Z-projected and analyzed with NIS-Elements AR Analysis software. For DeltaVision imaging, time-lapse images were Z-projected using the softWoRx software (Applied Precision, GE Healthcare). All images were processed using ImageJ for presentation.

#### Immunofluorescence

Cells were grown on Precision cover glass and fixed with 4% formaldehyde for 10 min, permeabilized with ice-cold methanol. Primary and secondary antibodies were diluted in PBS containing 5% bovine serum albumin (BSA) and incubated for 1-2 h. After antibody staining, samples were mounted on microscope slides with ProlongGold-DAPI (Invitrogen). Antibodies rabbit pericentrin 1:400 (Abcam, #ab4448); sheep α/β-tubulin 1:500 (Cytoskeleton Inc ATN02)

#### Transfections

Cells at 50% confluency were transfected with 5 nM siRNA using Lipofectamine RNAiMAX transfection reagent (Life Technologies) following the manufacturer’s instructions. GCN2 targeting siRNA sequence was gcaauucuguggugcauaa.

#### Cloning and lentiviral transductions

GCN2 constructs and mutants were as described [36]. Stable cell lines were generated by lentiviral transductions using procedures and plasmids that have been previously described [69]. For the experiments in Figures 6 E and F the destination vector pCW57.1 (Addgene, #41393) was used, which allows doxacycline-regulatable expression of the transgene. Detailed cloning procedures are available on request.

#### Migration assays

Wound healing assays were performed either by seeding cells into culture inserts (Ibidi, #80366) and observing them by live-cell microscopy after removing the insert, and images were analyzed using the NIS-Elements AR Analysis software (Fig 4D, Fig S3E). Alternatively, cells were grown to confluence in 96-well plates, wounded using the Sartoriusu Wound maker, observed them in an Incucyte, and images were analyzed with the Incucyte Scratch Wound Analysis Software (Fig 4C).

For transwell migration assays, cells were pre-incubated in FBS-free medium for 4 h, then seeded into cell culture inserts (Nunc, 8 µm pore size), still in FBS-free medium and exposed to an FBS gradient. After 16 h incubation, not-migrated cells in the chamber were removed, the migrated cells on the underside of the membrane were fixed, stained with DAPI and counted. Cell counts were normalized to the seeding controls, which were determined either by crystal violet staining and counting, or by the CCK cell proliferation assay (Abcam, #ab228554).

Chemotaxis assays (Fig 5, Fig S3D)) were performed using µ-Slide Chemotaxis slides (Ibidi # 80326), live-cell imaging images were analyzed and cells were tracked using the NIS-Elements AR Analysis software, and tracks were analyzed using the Chemotaxis and Migration Tool (Ibidi).

## Movies

Movie 1 SiHa cell going through mitosis in the absence of GCN2i. Related to Fig 3. Images were taken every 5 min, tubulin is shown in magenta, DNA is shown in gray.

Movie 2 SiHa cells delay mitotic progression in the presence of 2 µM GCN2i. Related to Fig 3. Images were taken every 5 min, tubulin is shown in magenta, DNA is shown in gray.

Movie 3 Caski cell going through mitosis in the absence of GCN2i. Related to Fig 3. Images were taken every 5 min, tubulin is shown in magenta, DNA is shown in gray.

Movie 4 Caski cells delay mitotic progression in the presence of 2 µM GCN2i. Related to Fig 3. Images were taken every 5 min, tubulin is shown in magenta, DNA is shown in gray.

Movie 5 Wound healing in control cells. Related to Fig 4E. Images were taken every 30 min.

Movie 6 Wound healing in cells incubated in the presence of 2 µM GCN2i. Related to Fig 4E. Images were taken every 30 min.

Movie 7 Wound healing in cells stably overexpressing GCN2 (GCN2^high^). Related to Fig 4E. Images were taken every 30 min.

## Notes

### Competing Interest Statement

The authors have declared no competing interest.

