## Supplementary figures and images for "Analysis of patient data reveals novel cancer-relevant functions for GCN2/eIF2αK4"

### Fig S1

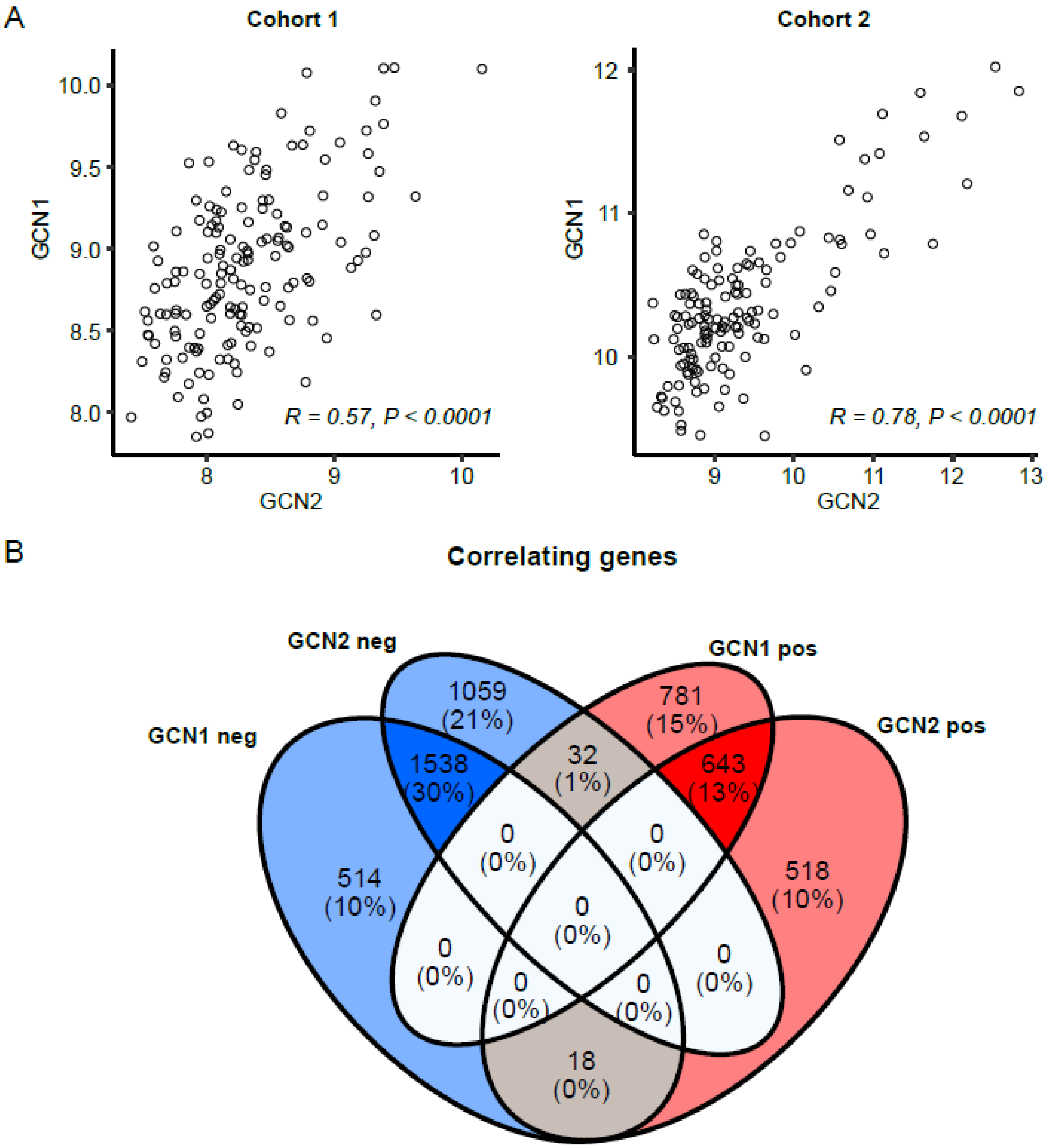

### Fig S3

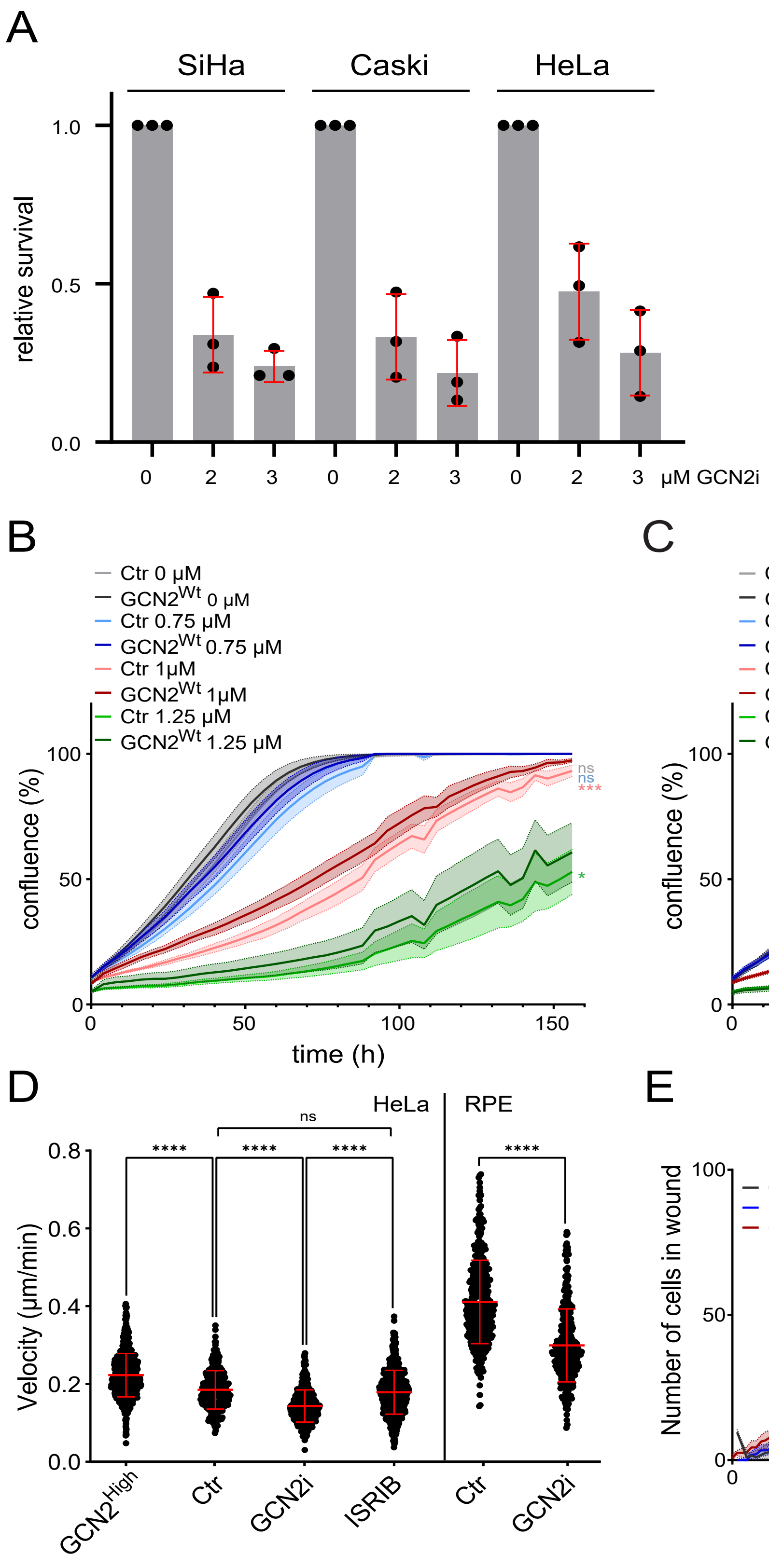

### Fig S4

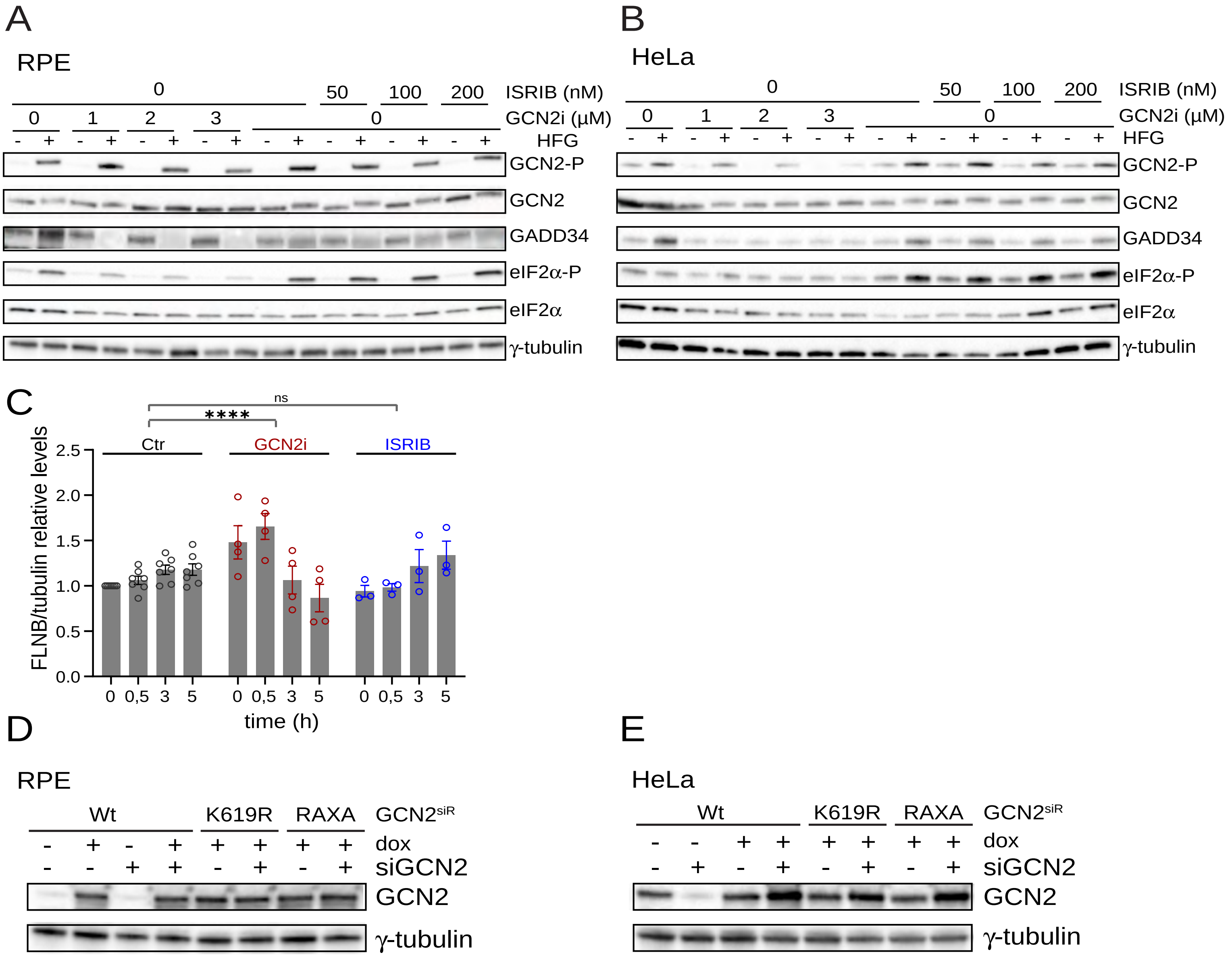
