## Supplementary material for "Analysis of patient data reveals novel cancer-relevant functions for GCN2/eIF2αK4": Fig S2

### Figure S2

## A

Enriched GO terms for genes overlapping with the hallmark “apical junction”

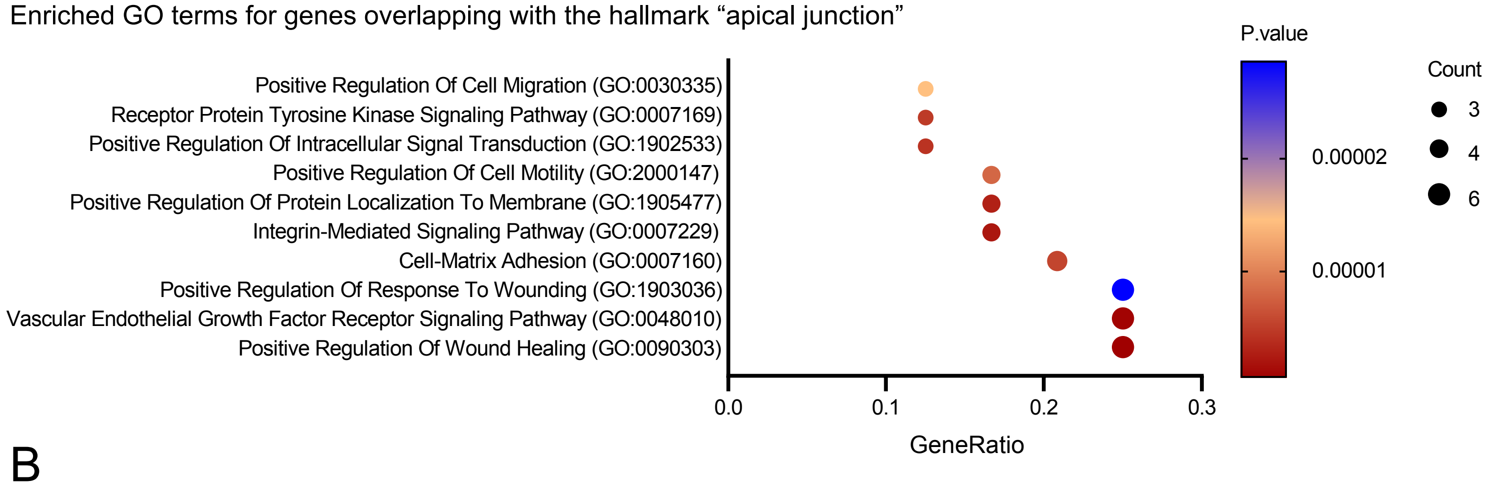


## B

Enriched GO terms for genes overlapping with the hallmark “mitotic spindle”

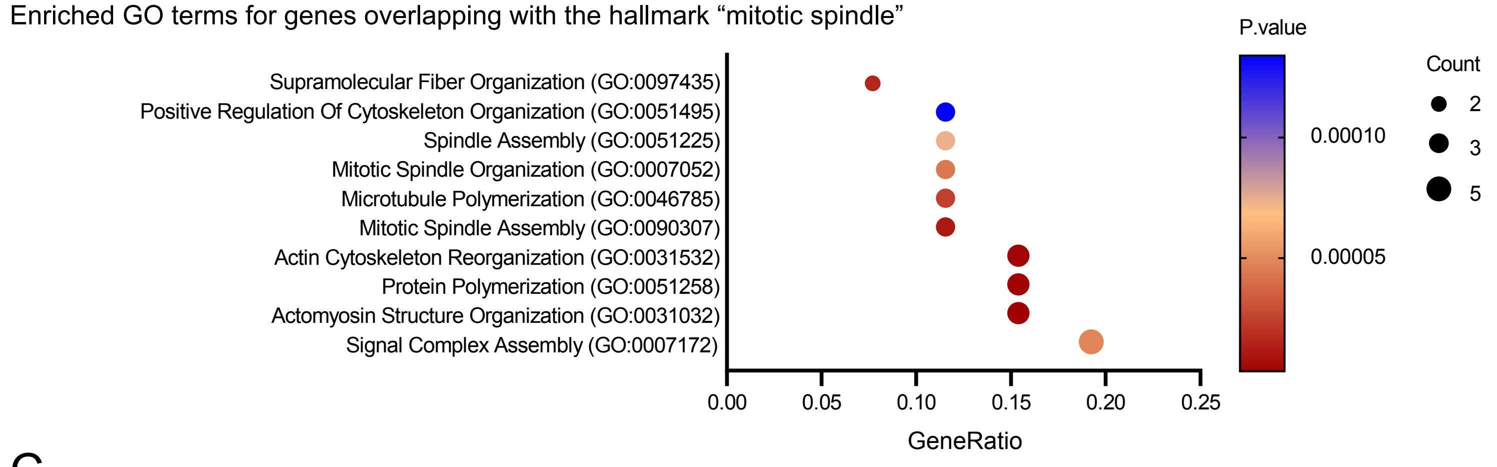


## C

Enriched GO terms for genes overlapping with the hallmark “G2/M checkpoint”

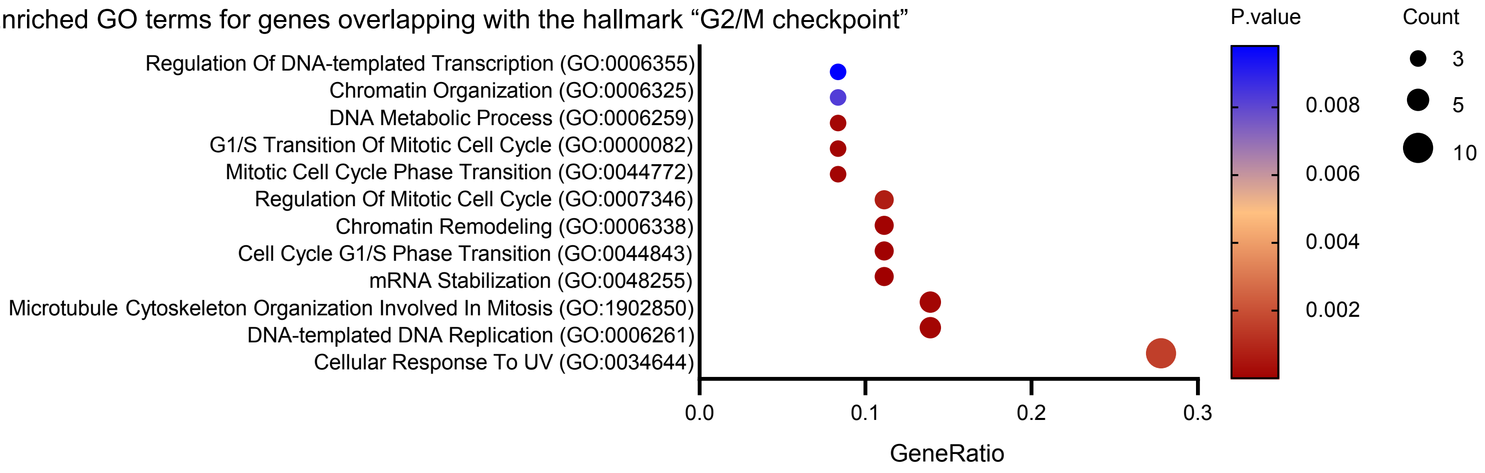
